# DNA Origami Signal Amplification in Lateral Flow Immunoassays

**DOI:** 10.1101/2024.07.05.602214

**Authors:** Heini Ijäs, Julian Trommler, Linh Nguyen, Stefan van Rest, Philipp C. Nickels, Tim Liedl, Maximilian J. Urban

## Abstract

Lateral flow immunoassays (LFIAs) enable a rapid detection of analytes in a simple, paper-based test format. Despite their multiple advantages, such as low cost and ease of use, their low sensitivity compared to laboratory-based testing limits their use in *e*.*g*. many critical point-of-care applications. Here, we present a DNA origami-based signal amplification technology for LFIAs. DNA origami is used as a structurally precise adapter to connect detection antibodies to tailored numbers of signal-generating labels. As a proof of concept, we apply the DNA origami signal amplification in a sandwich-based LFIA for the detection of cardiac troponin I (cTnI) in human serum. We show a 55-fold improvement of the assay sensitivity with 40 nm gold nanoparticle labels and an adjustable signal amplification of up to 125-fold with fluorescent dyes. The technology is compatible with a wide range of existing analytes, labels, and sample matrices, and presents a modular approach for improving the sensitivity and reliability of lateral flow testing.

---

Lateral flow immunoassays (LFIAs) are among the most widely used diagnostic tests worldwide. The scalable production, ease of use, and the possibility for rapid, instrument-free readout have made them a prime diagnostic tool in the medical field, particularly in the point of care (PoC). [1, 2]

While LFIAs can provide results in the PoC within minutes and with minimal effort, they are often limited by low sensitivity – *i*.*e*., detection limits ranging from micromolar down to 10–100 picomolar concentrations [3] – and thus the initial test result needs to be confirmed with additional laboratory-based tests. Standard laboratory-based immunoassays reach femto- to picomolar detection limits by using powerful signal amplification methods, such as enzymes (enzyme-linked immunosorbent assay; ELISA) and electrochemiluminescence (ECL), but they also require specialized equipment, stringent control over reaction conditions, and operation by trained personnel. [3, 4] Signal amplification strategies applicable directly in LFIAs hold great promise in improving their sensitivity and reliability at the PoC and in providing better rapid testing options for low-resource settings and home testing.

Various approaches to signal amplification on LFIAs have been demonstrated before. [5] The signal on commercial LFIAs is typically a colored line formed by antibody-conjugated optical labels, such as colloidal gold, where multiple antibodies are linked to a single label. [2, 6] As the signal strength correlates with the number of labels per detectable analyte molecule, this limits the signal intensity. To amplify the signal without changing the type of reporter, nanostructures with repetitive motifs have been applied to link antibodies to multiple labels. These polymerization approaches include the use of DNA dendrimers, hybridisation chain reaction, [7, 8] or DNA-tile assembly [9]. One drawback of these methods is that they allow only limited control over the size of the applied nanostructures and over the stoichiometry of antibodies and labels. Issues arising from polydispersity, amplification of unspecific signals, and aggregation of labels have so far hindered the integration of these approaches in commercial LFIA products.

DNA self-assembly allows the creation of nanostructures with Ångström precision on the scale of 10–1,000 nanometers. [10] In particular, DNA origami [11] is a robust molecular programming technique that has been used to arrange molecular binders including antibodies [12–18] and a wide variety of labels [19, 20]. Despite groundbreaking research work over almost two decades, DNA origami nanostructures are still not applied in mass produced products to improve human health or quality of life.

Here, we present a DNA origami-based signal amplification technology for lateral flow tests, and in particular, for LFIAs (Fig. 1). The technology relies on precise control over the stoichiometry of antibodies and labels. It is thus possible to detect one analyte with a DNA origami structure that, for example, carries exactly one detection antibody. This can already lead to an improved signal compared to conventional antibody-conjugated labels. Depending both on the label type and on the DNA origami design, the number of labels can range from one to up to several hundred. In addition to significantly amplifying the signal, the amplification factor can thus be adjusted according to meet the desired sensitivity of the signal-amplified LFIA.

**Fig. 1.**
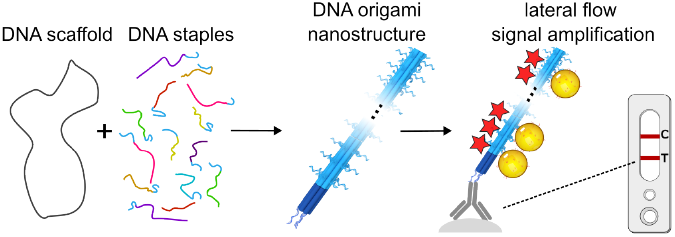
Lateral flow immunoassay signal amplification with DNA origami. Structurally precise DNA nanostructures are prepared with the DNA origami self-assembly method. The DNA origami is applied as an adapter that connects a specified, low number of detection antibodies to multiple labels. The greatly enhanced number of labels per antibody and per bound analyte at the test line leads to a stronger signal and makes lower analyte concentrations detectable by naked eye.

Our amplification technology is designed to be integrated to established LFIAs on the market, supporting compatibility with diverse analytes, detection molecules, as well as LFIA mass production. The technology can also be combined with the whole toolbox of novel labels that have been developed in the recent years [21–23]. Here, we focus on optimizing the technology using fluorescent dyes and 40-nm gold nanoparticles. As a proof of concept for the signal amplification, we demonstrate the detection of cardiac troponin I (cTnI) in human serum samples. cTnI is a key biomarker for cardiovascular diseases, playing a vital role in the urgent differential diagnosis of heart attacks, particularly in non-ST elevated acute coronary syndrome. [24, 25] LFIA tests for cTnI are globally utilized, especially in settings lacking central laboratory access. There remains a significant medical need for rapid (under 20 minutes) and sensitive (below 50 ng/l) cTnI tests at the PoC. As the current high-sensitivity cTnI assays have reached maturity through five generations of optimization, the cTnI test serves here as the established benchmark for assay development.

## The DNA origami signal amplification structure

The DNA origami signal amplification structure functions as an adapter that connects detection antibodies with labels in a tunable stoichiometry. Here, we have used a common six-helix bundle (6HB) design, which comprises six parallel, interconnected DNA duplexes, and has a length of 490 nm and a diameter of 8 nm in solution [26]. The main functional features necessary for signal amplification are the two binding domains at the ends of the 6HB and a signal amplification domain in between (Fig. 2a).

**Fig. 2.**
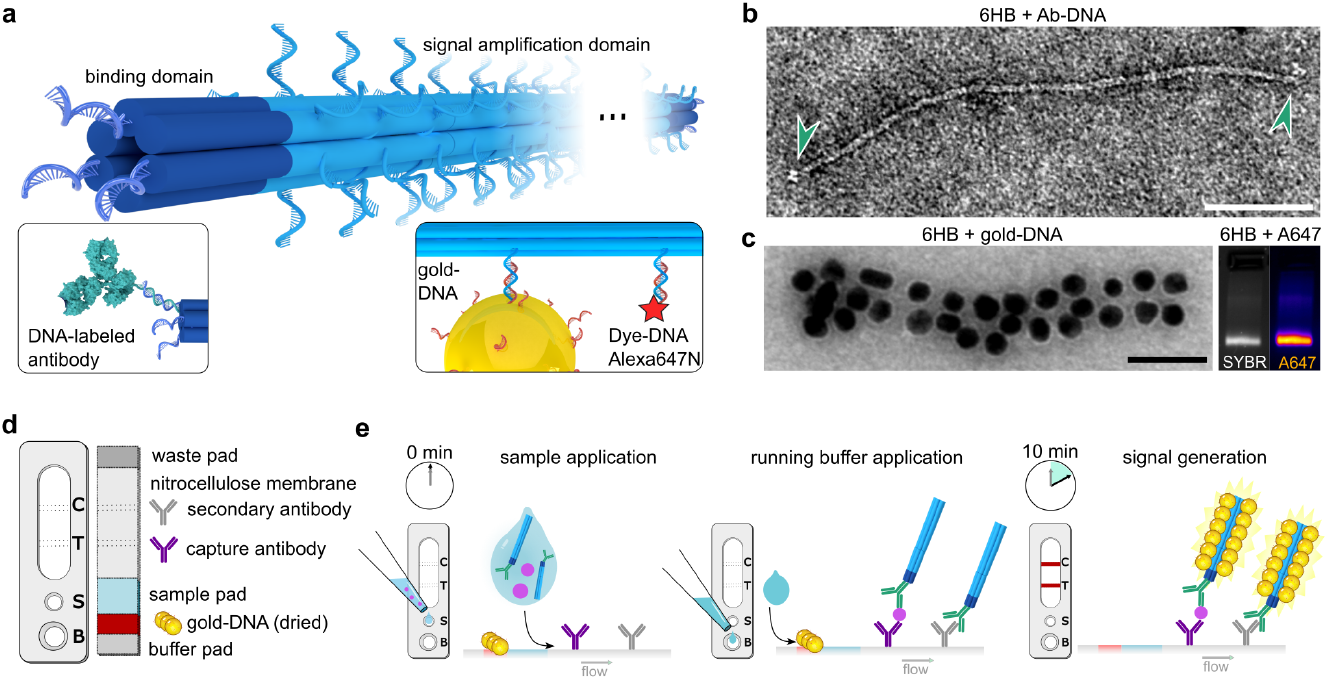
The DNA origami signal amplification method. **a**. A schematic presentation of the design of a DNA origami 6-helix bundle (6HB) with an adjustable number of binding domains (dark blue) and a signal amplification domain (light blue). For lateral flow immunoassays (LFIAs), antibodies are attached to the binding domains. DNA-conjugated labels, such as gold nanoparticles or fluorophores, bind to the signal amplification domain. **b**. Transmission electron microscopy (TEM) image of a detection antibody-6HB conjugate (Ab-6HB) after incubation of 6HBs with 6-fold molar excess of DNA-conjugated detection antibodies (Ab-DNA) and gel purification. The bound antibodies are indicated by arrows (scale: 100 nm). **c**. Decoration of the 6HBs with labels. The TEM image in the left panel shows a 6HB after incubation with DNA-labeled gold nanoparticles; here, 20 nm particles for the purpose of demonstration (scale: 100 nm). The right panel shows an agarose gel electrophoresis (AGE) analysis of Sybr Safe DNA stain and Alexa Fluor 647 (A647) fluorescence of 6HBs after incubation with a 200-fold molar excess of A647-labeled oligonucleotides. **d**. A general design of a LFIA test design used for the integration of DNA origami signal amplification. **e**. LFIA running protocol. The amplified signal is generated when the sample-DNA origami mixture interacts with both the labels and the immobilized antibodies on the test (T) and control (C) lines.

### Binding domains

The binding domains feature up to 12 single-stranded staple oligonucleotide (ssDNA) overhangs for either direct hybridization with oligonucleotide analytes or for attachment of DNA-conjugated recognition molecules, such as antibodies. In all experiments presented in this work, each binding domain has three overhangs, resulting in a total of six overhangs per 6HB. After an initial HPLC purification step to remove excess staple strands, the 6HBs were used for DNA target detection without further modification. For the detection of cTnI, the binding domains were functionalized with anti-cTnI IgG antibodies via DNA hybridization. The detection antibodies were conjugated to DNA handles using non-site-specific labeling of lysine residues (Fig. S1). The 6HBs were then incubated with a 6-fold molar excess of antibodies to form the Ab-6HB conjugates. The conjugates were employed in LFIAs without further purification. After characterizing the conjugates with agarose gel electrophoresis (AGE) (Fig. S2) and extracting them from the leading band, TEM revealed conjugated antibodies bound to the binding domains at the ends of the 6HBs (Fig. 2b and Fig. S3–S4). Typically, 1–2 antibodies per binding domain could be observed.

### Signal amplification domain

The signal amplification domain of the 6HB contains 180 ssDNA overhangs designed for binding DNA-conjugated labels. To demonstrate the signal amplification strategy, we used two different labels: DNA-functionalized 40 nm gold nanoparticles (gold-DNA) and Alexa Fluor 647-labeled DNA oligonucleotides (A647-DNA). The colorimetric signal of the gold-DNA labels is visible to the naked eye, while the fluorescence from A647-DNA can be detected using a fluorescence reader.

To study the attachment of gold-DNA labels to the 6HBs, the 6HBs were incubated with an excess of 20 nm gold-DNA labels, and subsequently gel purified. TEM analysis of the assemblies shows that the binding sites on the 6HB can facilitate a dense binding of labels along the entire length of the signal amplification domain (Fig. 2c, left panel). TEM images of the 6HBs complexed with 40 nm gold-DNA labels are shown in Fig. S5. The attachment of the A647-DNA labels to the 6HBs was studied by means of AGE. The presence of colocalized fluorescence signals from both the SYBR Safe DNA stain and the A647 confirms the hybridization of A647-DNA with 6HBs (Fig. 2c, right panel). Throughout this study, we used a 6HB featuring 3 ssDNA overhangs at each binding domain and 180 ssDNA overhangs at the amplification domain. The amplification factor was tuned by adjusting the molar ratio of A647-DNA or gold-DNA labels relative to the 6HB during the conjugation process (Fig. 2c and Fig. S6).

### DNA origami nanostructures on LFIAs

Signal amplification is achieved by incorporating the DNA origami structure into a conventional LFIA test strip comprising a nitrocellulose membrane striped with capture reagents – *e*.*g*., antibodies or streptavidin – as well as pads for sample application, buffer application, storage of dry reagents, and liquid absorption after the nitrocellulose membrane (Fig. 2d). For running assays with A647-DNA labels, the Ab-6HB conjugates were preassembled with the desired excess of A647 labels prior the assay. When utilizing gold-DNA, the labels were dried on a pad upstream of the sample application zone. The LFIA tests were then run in two steps (Fig. 2e). First, the liquid sample was mixed with the Ab-6HB conjugates and applied on the sample application pad. In the second step, running buffer was applied to release the dried gold-DNA labels. This induces a capillary flow that carries all components to the test and control lines, where they can interact with the capture reagents. An intense colorimetric signal is generated when a large number of gold-DNA labels bind to the signal amplification domains of the 6HBs. Drying of the particles is not only imperative for the long-term stability of the gold labels and for maintaining a simple user protocol without additional label handling, but it also leads to an improved test performance through a slower release of the gold-DNA labels. This was observed to prevent unwanted early interactions that can cause label-mediated crosslinking, sedimentation of high-molecular-weight 6HB-gold assemblies, and signal loss at the test line.

## Controlling the amplification factor

To show that the DNA origami allows for a precise control over the signal amplification factor on lateral flow tests, we used the 6HBs with A647-DNA labels for detecting a biotinylated DNA oligonucleotide. By selecting biotin-DNA as the model analyte, the amplification system can be studied without limitations arising from *e*.*g*. potential slow or low-affinity antibody-antigen binding reactions. The biotin-DNA can be detected without antibodies through two fast (high forward rate constant, *k*_*on*_) and high affinity (low dissociation constant, *K*_*d*_) binding reactions: biotin-streptavidin interaction (*K*_*d*_ = ∼10^−15^ mol/l) and the hybridization of a 26 base pairs long DNA double strand. The illustration in the left panel of Fig. 3a shows the molecular schematic of the structure that forms at the test line in the assay. When a sample containing the complementary biotin-DNA analyte is mixed with the A647-labeled 6HBs, the analyte is bound through DNA hybridization at the binding domains. The 6HB-bound analytes are then captured at a streptavidin test line.

**Fig. 3.**
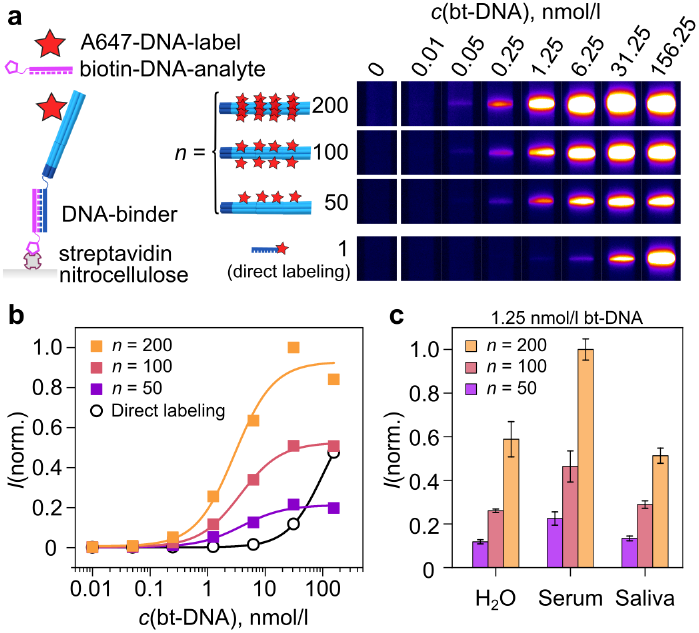
Proof-of-principle detection of biotinylated DNA and adjustable fluorescence signal on lateral flow assays. **a**. Detection of a biotin-DNA analyte (26-mer DNA oligonucleotide with a 3’ biotin) in water. The 6HBs used in the lateral flow assay were mixed with either 50-, 100-, or 200-fold excess (*n*) of A647-DNA labels per 6HB prior use. The non-amplified (*n* = 1) reference (bottom panel) contains no 6HB, but instead A647-DNA that is complementary to the biotin-DNA analyte.The images show the A647 fluorescence at the test line after 15 min assay time, and they have all been captured using a 16 ms exposure time. **b**. Normalized fluorescence intensities of the test line images shown in Fig. a. Each set of intensity data has been fitted with a four-parameter logistic function. **c**. Normalized test line intensities for 1.25 nmol/l biotin-DNA in water, human serum, and saliva, detected with 6HBs with different amounts of A647-DNA. The results are presented as the mean *±* standard deviation of three repeated experiments.

Prior running the biotin-DNA fluorescence assays, 6HBs were mixed with either 50, 100, or 200 A647-DNA labels per 6HB (*n* = 50, 100, or 200) to adjust the amplification factor. A small volume (5 *µ*l) of biotin-DNA in milli-Q H_2_O was mixed with the 6HB-A647, added on the test strip, and flushed over the nitrocellulose membrane with running buffer. As an unamplified reference (*n* = 1; bottom row of Fig. 3a), the biotin-DNA analyte was mixed with a complementary, A647-labeled DNA oligonucleotide instead of 6HB-A647. The concentration of the direct labeling probe was the same as the concentration of A647-DNA labels in the *n* = 100 sample.

The test line fluorescence images presented in Fig. 3a and the fluorescence intensity values plotted against the biotin-DNA analyte concentration in Fig. 3b show that the fluorescence intensity can be both amplified and controlled with the 6HBs. In the presented experiment, only 6.25 nmol/l or higher concentrations of biotin-DNA were detectable with direct labeling (*n* = 1). At smaller concentrations, the number of fluorophores at the test line is below the threshold that can be distinguished from the background fluorescence. 6HBs increase the number of fluorophores at the test line, and thus the sensitivity of the assay increases with *n*. With *n* = 200, biotin-DNA concentrations down to 50 pmol/l could be detected, corresponding to ∼125 -fold higher sensitivity than in the case of *n* = 1. The sensitivity improvement by two orders of magnitude is well in line with the expected signal amplification factor; 200 A647-DNA labels per 6HB can be expected to saturate all of the 180 label binding ssDNA overhangs of the signal amplification domain and yield the maximum 180-fold signal amplification achievable with the applied 6HB design. Moreover, Fig. 3b shows that the average fluorescence signal from the 6HBs decreases by 41% when *n* is decreased from 200 to 100, and by further 66% when going from *n* = 100 to 50, thus closely following the excess of labels used in the self-assembly reaction.

While the higher label-analyte stoichiometry is the main determining factor for the increased signal and sensitivity at low and mid-range biotin-DNA concentrations, the effects of other experimental factors increase at the highest concentrations. The saturation intensities for each *n*, for instance, depend on the choice of experimental parameters, such as the amount of 6HBs used per strip, the number of analyte binding sites per 6HB, the applied running protocol, and the capacity of the test line. By adjusting the aforementioned parameters, the response curves of the 6HB signal amplification system can be tuned to fit the most important concentration range of the selected analyte.

Finally, we verified that the signal amplification shown in Fig. 2a–b for biotin-DNA can be reproduced in complex sample matrices. To show this, we compared the detection of biotin-DNA in water, human serum, and human saliva. The lateral flow assays were again run by mixing the sample with the A647-labeled 6HBs before application on the strip, which in this case has the importance of bringing the 6HBs into a direct contact with each sample matrix before dilution by the running buffer takes place on the test strip. As seen in Fig. 2c, the 6HBs were compatible with all tested sample matrices and the signal amplification factor could be controlled with *n*, with a slight variation of the fluorescence intensities depending on the type of sample. Blank samples without biotin-DNA showed no signal nor nonspecific binding in any of the studied sample matrices (Fig. S7).

## Detection of cardiac troponin I

We then constructed a full antibody sandwich-based LFIA for the detection of cTnI. To achieve this, capture antibodies and detection antibodies need to be introduced into the assay. Here, we used the same lateral flow test strips with a streptavidin test line as for validating our method with biotin-DNA and a combination of biotinylated capture antibodies (Ab_capture_-biotin) and detection antibody-6HB conjugates (Ab_1_-6HB or Ab_2_-6HB). In addition to detecting cTnI in the A647 fluorescence assay, we combined the 6HB signal amplification with gold-DNA labels for a higher sensitivity and an instrument-free readout.

To study the sensitivity and specificity of the cTnI detection in a real sample matrix, we used serum samples with known cTnI concentrations. For this, we spiked human serum with recombinant cardiac troponin IC complex (cTnIC). cTnIC is one of the three forms of cTnI present in the circulation after an acute myocardial infarction – free cTnI, the binary cTnIC complex, and the ternary complex of troponin I, C, and T.[27] The higher stability of cTnI in the IC complex than in the free form makes cTnIC a reliable concentration standard in the assay development. Despite the presence of troponin C, all serum troponin concentrations are reported as the concentration of cTnI.

The principle of the cTnI detection and the A647 fluorescence assay is described schematically in the left panel of Fig. 4a. To run the assays, a small volume (5 *µ*l) of human serum was first mixed with both the Ab_1_-6HB conjugate and the Ab_capture_-biotin. The mixture was applied on the sample application pad of the test strip and flushed over the nitrocellulose membrane with running buffer. As an unamplified reference (*n* = 1), the DNA handles on the Ab_1_-DNA were labeled directly with a complementary A647-DNA oligonucleotide. The smallest detectable concentration with direct labeling was 1.25 nmol/l (Fig. 4a). When using Ab_1_-6HB with *n* = 100, fluorescence at the test line could be detected down to 10 pmol/l concentration of cTnI. This corresponds to *ca*. 100-fold improvement of the LoD. A consistent 100-fold higher A647 fluorescence intensity was observed for all the cTnI concentrations where the unamplified signal is available for comparison. Full LFIA strip images are presented in Fig. S8 and show that comparable signal amplification takes place also at the control line.

**Fig. 4.**
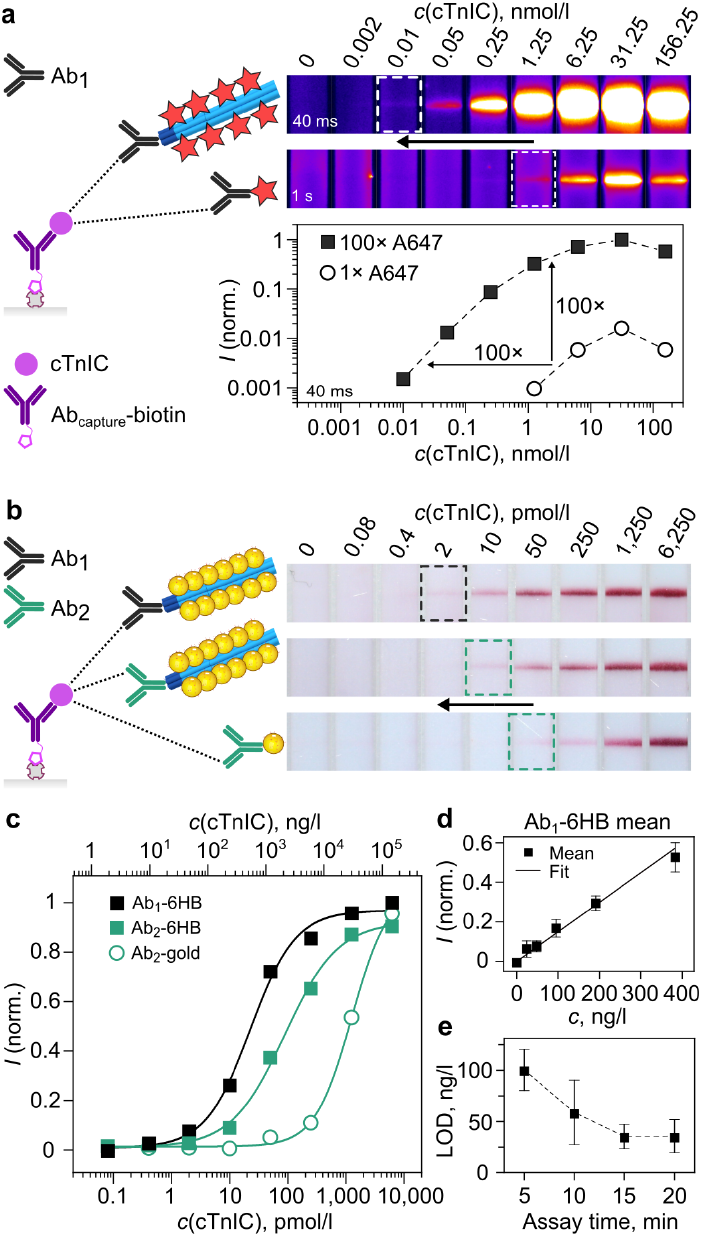
LFIA detection of cardiac troponin IC complex (cTnIC) in human serum. **a**. A647 fluorescence assay. The left panel describes the assembly of the 6HB signal amplification complex at the test line. The right panel shows test line fluorescence images after 15 min assay time when using either Ab_1_-6HBs with 100 A647-DNA labels, or Ab_1_-DNA conjugate that is directly labeled with a complementary A647-oligonucleotide. For higher visual clarity, test line images captured using a longer exposure time of 1 s are shown for direct labeling. Visual limits of detection (LoD) of both data sets are highlighted. Test line intensities captured with 40 ms exposure are plotted against the cTnIC concentration in the bottom panel. **b**. Gold-DNA assay with Ab_1_-6HBs, Ab_2_-6HBs, and state-of-the-art Ab_2_-gold conjugates. The photographs in the right panel show the test lines of assay strips after 15 min running time. The same capture antibody (Abcapture-biotin) was used in all experiments. The visual LoD of each data set is highlighted. **c**. Binding curves constructed from the 15 min test line intensities of Fig. b. All data sets have been fitted with a 4-parameter logistic function. **d**. Test line intensities near the LoD for Ab_1_-6HB, displayed as a mean *±* standard deviation of three repeated experiments. **e**. Dependency of the LoD on the assay time for Ab_1_-6HB.

To run the 6HB assays with gold-DNA labels, we used test strips with dried gold-DNA labels (Fig. 2d) and a running protocol described in Fig. 2e, with the exception that the serum samples were mixed with both the Ab-6HB and the Ab_capture_-biotin before the assay. While the same Ab_capture_-biotin was used on all test strips, here we compared the performance of two Ab-6HB conjugates: Ab_1_-6HB and Ab_2_-6HB. A broader screening of different antibody pairings with the A647 assay is shown in Fig. S9. As a reference for state-of-the-art sensitivity, we prepared Ab_2_-gold labels using a passive adsorption protocol. For LFIAs, a 5-*µ*l serum sample was then mixed with the Ab_2_-gold and the Ab_capture_-biotin before application on the test strip. The amount of gold particles (gold-DNA or Ab_2_-gold) was identical on all strips.

Fig. 4b shows a comparison of the cTnI detection sensitivity of the 6HBs with gold-DNA labels and the Ab_2_-gold labels. In the presented 1:5 dilution series of cTnIC in serum, the visual LoD of 50 pmol/l for Ab_2_-gold could be decreased to 10 pmol/l with Ab_2_-6HB, a 5-fold sensitivity increase. By optimizing the antibody sandwich with Ab_1_-6HB, the visual LoD could be decreased down to 2 pmol/l, corresponding to a further 5-fold sensitivity increase. A more detailed analysis of the test line intensities in Fig. 4c shows that a 14-fold improvement in assay sensitivity was achieved with the Ab_2_-6HB signal amplification, and an overall amplification factor of 55 with Ab_1_-6HB. While Ab_1_-gold conjugates would undoubtedly also improve the sensitivity of the state-of-the-art detection, the production of Ab_1_-gold particles with the passive adsorption protocol established for Ab_2_-gold was unsuccessful. Interestingly, both Ab_1_ and Ab_2_ could still be readily labeled with DNA and conjugated to the 6HBs. This shows the robustness of the Ab-6HB conjugation method even with challenging antibodies that are, for example, prone to self-association.

Comparison of the signal amplification outcome between the A647-DNA and the gold-DNA labels (Fig. 4a–b) highlights the characteristics of each type of label. In line with the label attachment on the 6HBs shown in Fig. 2c, the signal amplification factor follows the physical constraints of each type of label. The small size of A647-DNA means that the number of labels per origami can be precisely controlled over two orders of magnitude (1–180 labels per 6HB). This was confirmed in the experiments shown in both Fig. 3 and Fig. 4a. The larger 40 nm gold-DNA labels can be attached to the 6HBs in lower numbers and likewise, a signal amplification of one order of magnitude was observed (Fig. 4b–c). Even with a lower number of labels per 6HB, the gold-DNA labels provide a high contrast through their high extinction and in our experiments, they outperformed the A647-DNA labels in terms of sensitivity.

A further study of the LoD of the Ab_1_-6HB detection was carried out by detecting cTnI with the gold-DNA labels in the low concentration region (1–64 pmol/l in serum) in three repeated experiments. Fig. 4d shows the linearity of the test line signal in this region after 15 min assay time. The corresponding data for 5, 10, and 20 min assay times are presented in Fig. S10. As shown in Fig. 4e, the LoD determined from the data decreases with assay time. This happens both as more gold-DNA labels bind to the 6HBs and increase the test line signal, and as unbound labels migrate to the liquid absorption (waste) pad and improve the contrast between the test line and the nitrocellulose membrane. Here, a final LoD of 1.5± 0.5 pmol/l (corresponding to 35± 12 ng/l) was reached in 15 minutes. It should be noted that the optimal assay time and the LoD are strongly dependent on experimental parameters, such as the test strip materials and size, the arrangement of sample and conjugate pads, the sample volume, and the volume and the composition of the running buffer. In this study, we have not focused on optimizing these parameters for the 6HBs. Put together, our results show that under the applied conditions, the 6HB signal amplification increases the assay sensitivity and retains the critical short assay time (*<* 20 min) despite the more complex assembly of assay components on the test line.

## Discussion

DNA origami allows for the programmable self-assembly of megadalton-scale structures with precisely controlled size, shape and addressability. Due to this exceptional level of control over shape and molecular function, DNA origami structures have established their role in academic research as advanced nanoscale measurement tools, addressable devices, and structural scaffolds. [5, 14, 28, 29] An essential question remaining within the DNA nanotechnology field is, however, how these unique advantages can be harvested to overcome current technological limitations and provide value in everyday life or commercial applications.

Our signal amplification method presents a general strategy for the development of new LFIA assays. Its modularity is based on the ease of arranging organic molecules and inorganic particles on DNA origami at predefined sites and in precisely controlled amounts. In state-of-the-art LFIAs, the stoichiometry of antibodies to labels is always larger or equal to 1. For 40 nm gold with passively adsorbed antibodies, dozens of antibodies are typically bound to a single label. [30] Moreover, the number of detection antibodies often scales with the total surface area of the labels, *i*.*e*., it increases with the amount or size of the labels. [6] In our case, the amount of antibodies and of label entities per test can be adjusted separate of each other. Here, we have used 1–6 antibodies linked to up to 180 dye molecules or up to 25 40 nm gold labels to boost the amplification factor and thus the overall LFIA sensitivity. Generally, adjusting the sensitivity of state-of-the-art assays can be laborious and often requires revisiting the initial antibody selection. Tuning the ratio between labels and detection molecules, in contrast, allows for simple sensitivity tuning.

A second major benefit of DNA origami relies on its roots in molecular programming. While other methods of nanofabrication always lead to a distribution of sizes and shapes, DNA origami structures are inherently exact macromolecular copies of each other. In the application on LFIAs, this excellent size and shape control of the DNA origami conjugates leads to well-defined migration and fast diffusion within the porous nitrocellulose membrane compared to heterogeneously-sized labels obtained in uncontrolled polymerization approaches. With its length of 490 nm and width of 8 nm, our DNA origami 6HB can be considered a flexible rod, possibly allowing for a reptating, unhindered motion through the porous matrix [31]. As both the DNA origami and the 40 nm gold particles are small compared to systems that rely on direct increase of label size, our platform profits from the high diffusivity of its components, which in turn can lead to increased LFIA sensitivity. [6]

Next to mere assay performance, it is important to consider the manufacturing costs for DNA origami-based LFIAs. To be widely available, LFIA tests need to be mass-produced at low costs. Consequently, signal amplification technologies aimed for this platform should not increase the costs of materials and production. In our experiments, we applied 12 fmol (77 ng) of DNA origami to each LFIA test. At a cost of 350 EUR per nmol of DNA origami (phosphoramidte solid-phase oligonucleotide synthesis, in-house produced p8634 scaffold) this amounts to an increase of less than one cent of material costs per test. Moreover, in commercial LFIA tests, the detection antibodies are a major source of material costs. As our DNA origami adapter removes the strong link between number of antibodies and number of labels, it also allows minimizing the amount of antibodies per test, potentially leading to reduced overall costs of the signal-amplified LFIAs.

In conclusion, we have shown that DNA origami can be integrated into LFIAs for simple, adjustable, and user-friendly signal amplification for the detection of analytes in relevant sample matrices such as water, saliva, and serum. Using cTnI as a benchmark, we performed a direct sandwich-based assay, where the number of labels per cTnI was increased to achieve signal amplification factors of approximately 100 for fluorescent dyes and 14 for gold nanoparticles. A LoD of 35 ng/l – or 1.5 pmol/l – was reached in an assay time of 15 minutes with gold labels, complying with the need for fast, sensitive, and simple PoC test devices. When compared to existing cTnI assays, the obtained sensitivity lies between the low sensitivity of conventional cTnI LFIAs on the market – comparable to our state-of-the-art gold-antibody conjugates – and the high-sensitivity cTnI laboratory assays with typical LoDs of *<* 10 ng/l [25]. In addition to the full-length IgG antibodies used here, detection molecules such as aptamers, engineered antibody fragments, and oligonucleotides can be linked to DNA origami structures via established conjugation approaches. In addition, our method allows for the simultaneous use of multiple labels in multiplexing assays. Our work presents a step towards the integration of DNA origami nanostructures into commercial, everyday products. Due to the versatility of the approach, it has broad application potential for the development of diagnostic tools and it will enable expanding the range of analytes detectable by conventional LFIA testing.

## Declarations

- Conflict of interest. The authors declare the following competing financial interest: A patent application (PCT/EP2022/065889) has been submitted.

