## Supporting Information for "DNA Origami Signal Amplification in Lateral Flow Immunoassays"

### Supporting Information for
### DNA Origami Signal Amplification in Lateral Flow Immunoassays

Heini Ijäs *, Julian Trommler, Linh Nguyen, Stefan van Rest, Philipp C. Nickels, Tim Liedl, Maximilian J. Urban *

Faculty of Physics and Center for NanoScience (CeNS), Ludwig-Maximilians-University, Geschwister-Scholl-Platz 1, Munich, 80539, Germany.

### Supplementary figures

#### Fig. S1: Supplementary data of antibody-DNA conjugation

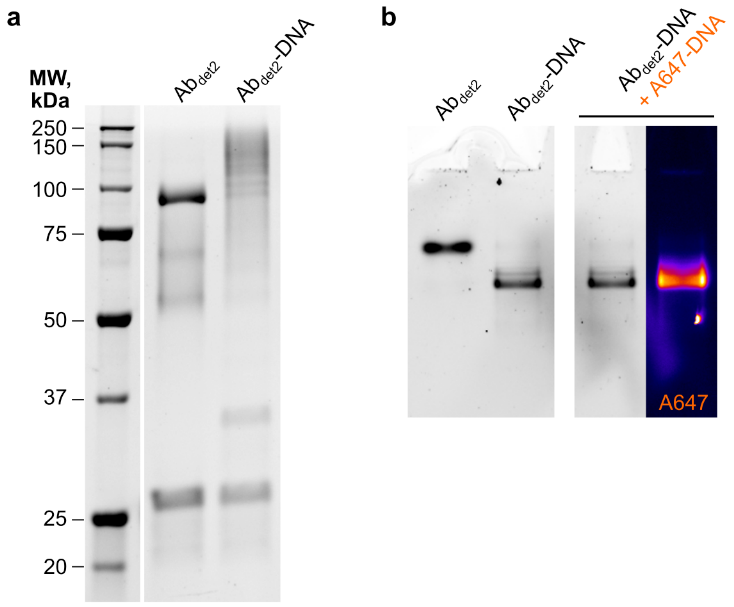

**Fig. 1** Gel analysis of antibody-DNA conjugation. **a**. SDS-PAGE analysis of the outcome of labeling $Ab_{det2}$ (anti-cTnI 19C7cc) with DNA. Left lane ($Ab_{det2}$) shows unmodified antibody. $Ab_{det2}$-DNA is the labeling product with a degree of labeling (DoL = $n$(DNA):$n$(Ab) of 2.85. Under denaturing conditions, the DNA-conjugated protein domains (IgG heavy and light chains) migrate slower than in the unmodified state due to the higher molecular weight. **b**. Native PAGE analysis of the DNA labeling of $Ab_{det2}$. The first two lanes contain the same samples as in Fig. a. In the third lane, the DNA-labeled antibody has been mixed with a 10-fold molar excess of the A647-DNA direct labeling probe. Under native conditions, the higher negative charge of DNA-labeled antibodies leads to higher electrophoretic mobility. Because native PAGE preserves the non-covalent interactions and thus both protein folding and DNA hybridization, the binding of the A647-DNA to the DNA handles on the antibody in the third lane can be observed through colocalization of signal at the protein stain channel (left) and the A647 fluorescence channel (right).

#### Fig. S2: AGE analysis of antibody-6HB conjugate formation

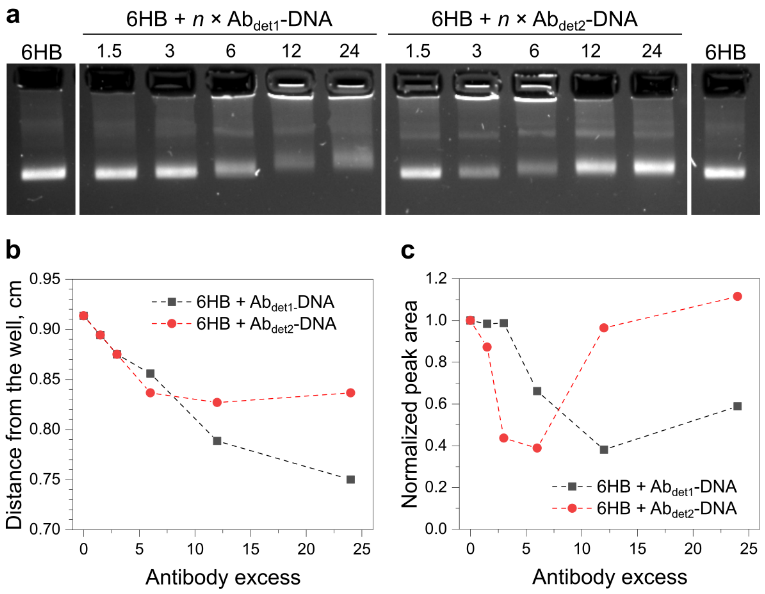

**Fig. 2** Characterization of the formation of $Ab_{det}$-6HB conjugates with the detection antibodies $Ab_{det1}$ and $Ab_{det2}$. **a**. AGE analysis of $Ab_{det}$-6HB mixtures prepared with different $Ab_{det}$:6HB molar ratios. The reference 6HB samples have been prepared without antibodies in the same buffer conditions as the $Ab_{det}$-6HB samples. $Ab_{det1}$-DNA is a DNA-conjugated rabbit anti-cTnI antibody Y302 with a degree of labeling of the DNA conjugation (DoL $= n(\mathrm{DNA}){:}n(\mathrm{Ab})$) of 0.94 determined from the UV-Vis absorbance spectrum. $Ab_{det2}$-DNA is a DNA-conjugated mouse anti-cTnI antibody 19C7cc with a DoL of 2.85. **b**. The electrophoretic mobility shift of the leading band of samples in Fig. a. **c**. The intensity of the leading band in the samples in Fig. a.

#### Fig. S3–S4: Additional TEM images of antibody-6HB conjugates

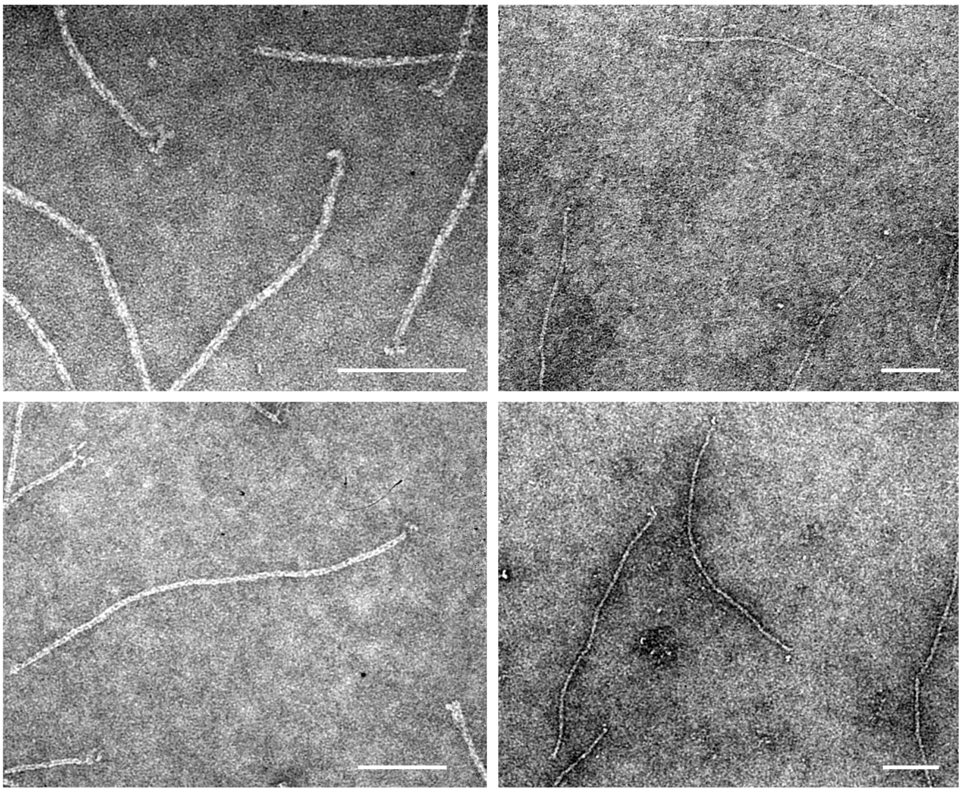

**Fig. 3** TEM images of $Ab_{det1}$-6HB conjugates after incubation of 6HBs with a 6-fold excess of $Ab_{det1}$-DNA and gel purification. All scale bars: 100 nm.

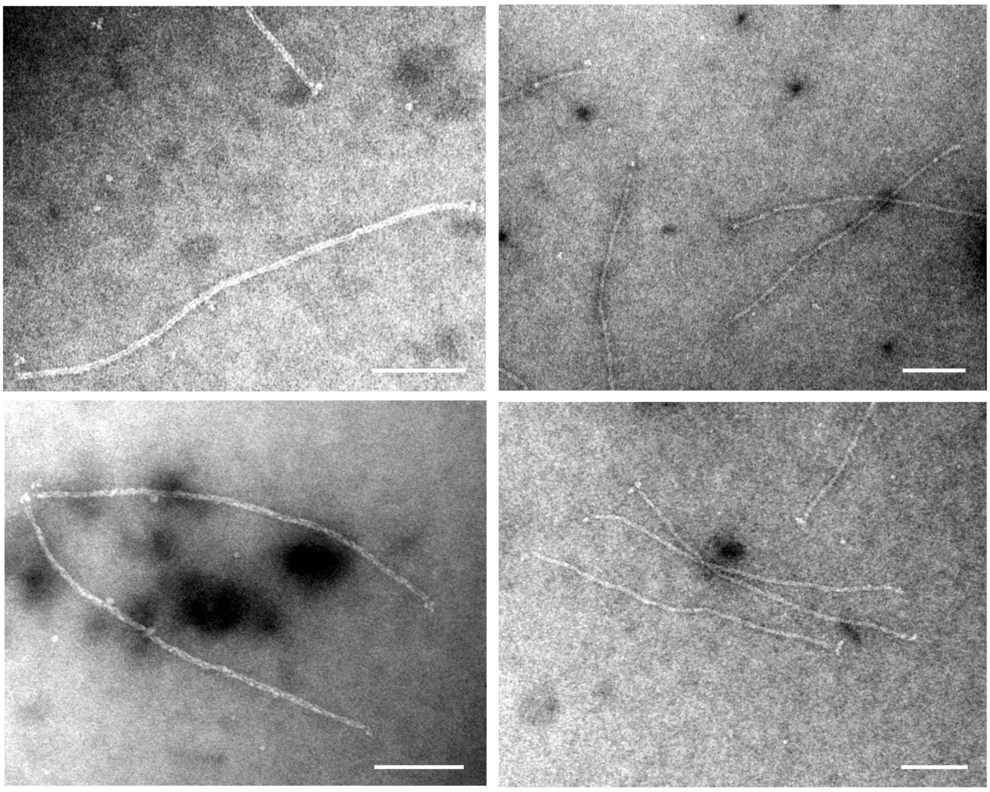

**Fig. 4** TEM images of unpurified $Ab_{det1}$-6HB conjugates after incubation of 6HBs with a 6-fold excess of $Ab_{det1}$-DNA without gel purification. Note that a fraction of antibodies can contain two or more DNA labels and lead to occasional coupling of two 6HBs, as visualized in the bottom left panel. All scale bars: 100 nm.

#### Fig. S5: Additional TEM images of 6HBs with 40 nm gold-DNA labels

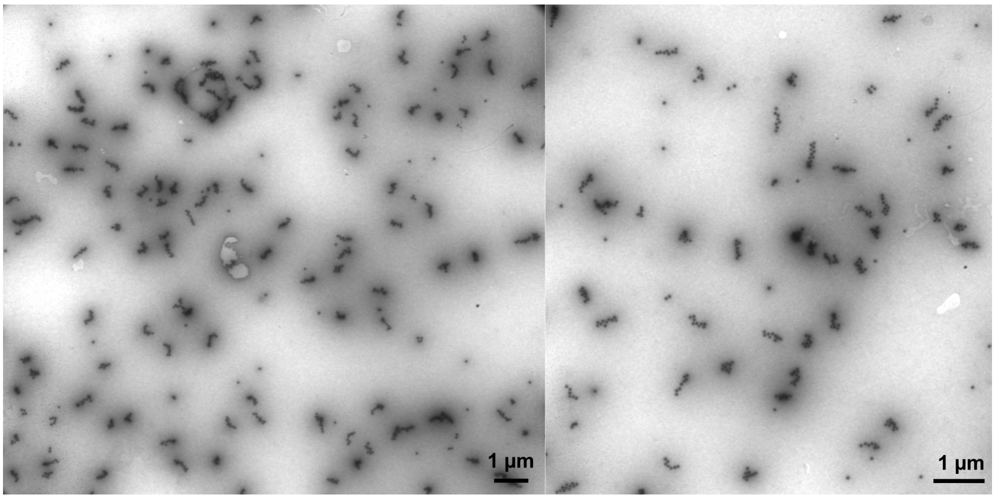

**Fig. 5** TEM images of gel-purified 6HBs with 40 nm gold-DNA labels.

#### Fig. S6: AGE analysis of the attachment of A647-DNA labels on 6HBs

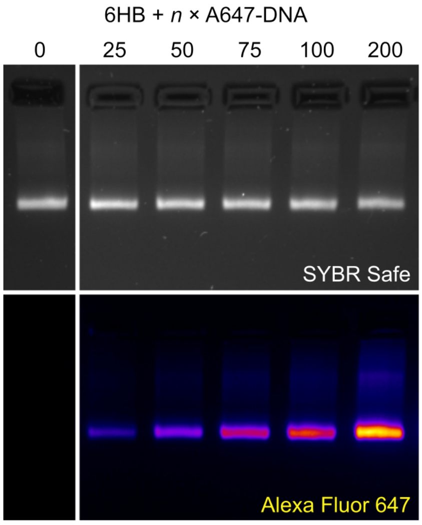

**Fig. 6** AGE characterization of mixtures of 6HBs and A647-DNA labels with different A647-DNA:6HB molar ratios. Both the fluorescence of the SYBR Safe DNA stain (top panel) and the A647 (bottom panel) are shown. The mixtures of A647-DNA labels and 6HBs were incubated at RT for ca. 20 minutes before the gel analysis.

#### Fig. S7: Fluorescence strip images with 0 nmol/l biotin-DNA in water, serum, and saliva

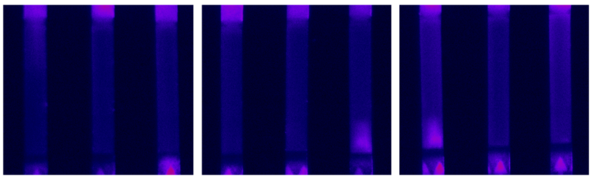

**Fig. 7** A647 assay strips with different sample matrices without analyte in the biotin-DNA detection experiments. To run the strips, 6HBs were mixed with a 200-fold excess of A647-DNA labels before the assay for a maximum signal. 5 $\mu$L of either milli-Q $H_2O$, human serum, or human saliva was mixed with the 6HBs, applied on the test strip after a short (ca. 1 min) incubation, and flushed over the nitrocellulose membrane with running buffer. The strips were imaged with a ChemiDoc MP fluorescence gel imager in the Alexa Fluor 647 channel after 15 min running time. For each sample matrix, three repeats are shown.

#### Fig. S8: Full LFIA strip images from cTnI fluorescence assays

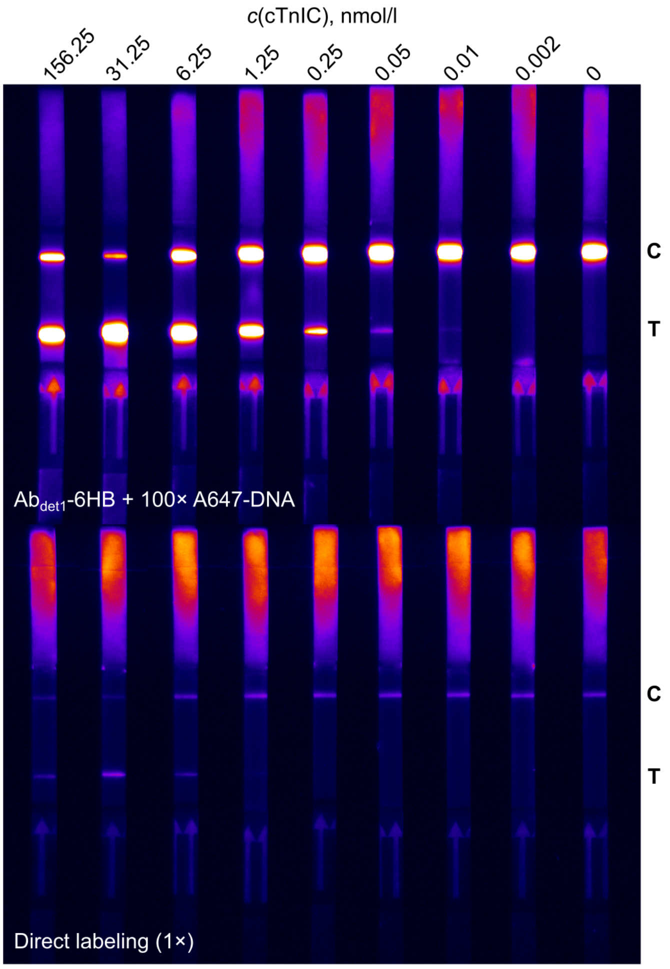

**Fig. 8** Full strip images of the cTnI A647-DNA assay presented in main text Fig. 4a. All images have been captured using a 40 ms exposure time and show the assay result after 15 min run time. The locations of the control (C) and test (T) lines on the membrane are indicated.

### Fig. S9: Screening of antibody pairings for cTnI detection with the fluorescence assay

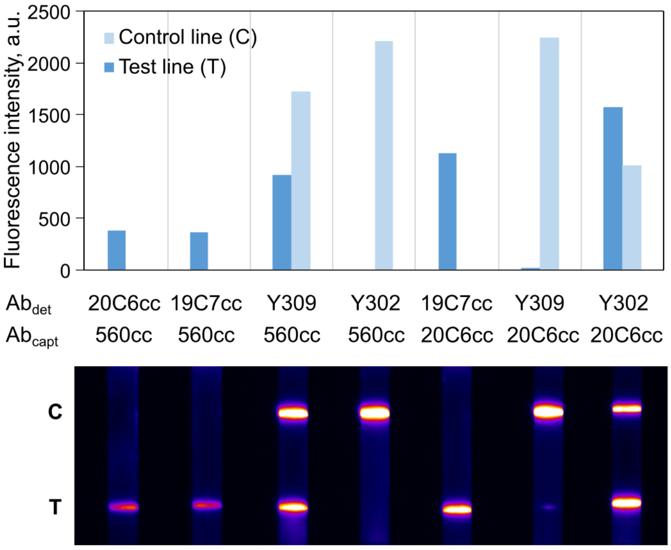

**Fig. 9** cTnI detection with different antibody combinations in the A647 6HB assay. The top panel shows test and control line intensities with the antibody combinations listed in the middle panel when detecting 1 nmol/l cTnIC in serum. The bottom panel shows the A647 fluorescence on the test strips after 10 min assay time. The LFIAs were run using the protocol presented in the main text. $Ab_{det}$ refers to the DNA-labeled detection antibody in the antibody-6HB conjugate, and $Ab_{capt}$ to the biotinylated capture antibody. Antibody-6HB conjugates were formed by mixing 6HBs with a 6-fold molar excess of different DNA-labeled $Ab_{det}$. Out of the presented $Ab_{det}$, Y309 and Y302 are rabbit IgG antibodies and thus captured at the control line by an anti-rabbit antibody, all other antibodies are mouse IgG antibodies.

#### Fig. S10: Supplementary data for cTnI LoD determination

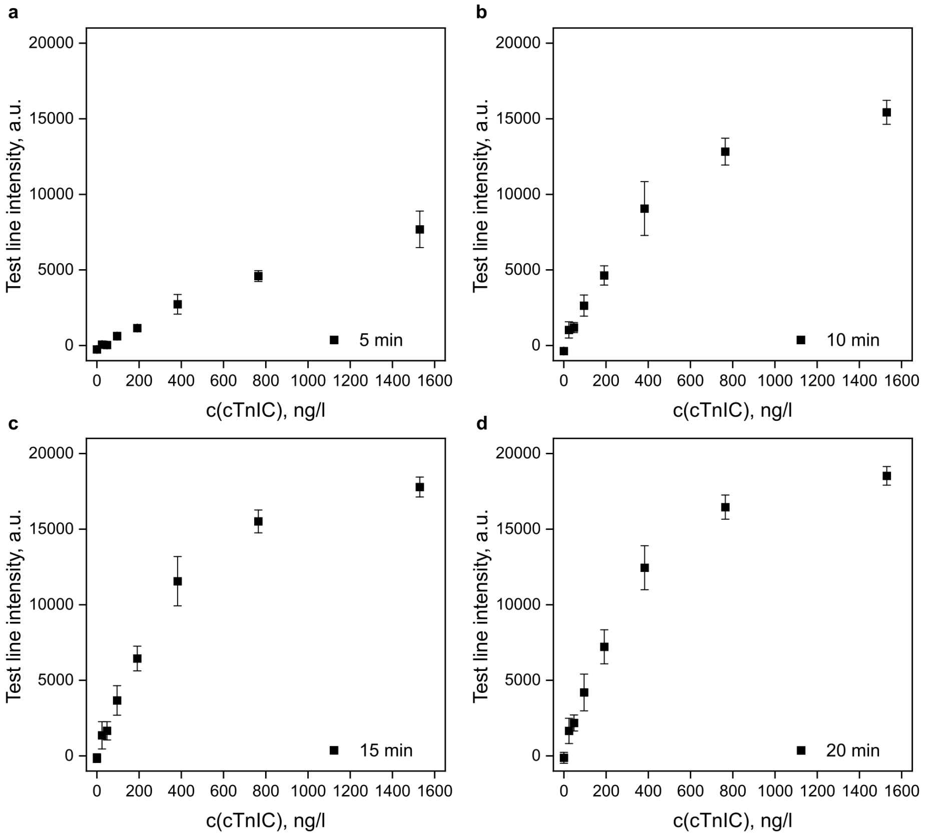

**Fig. 10** Test line intensities after different assay times used for determining the LoD of cTnI detection with $Ab_1$-6HB and gold-DNA labels. All data are shown as the average of three repeated experiments ± standard deviation. For all data setes, intensity values between 24–382 ng/l (1–16 pmol/l) were termed the linear response region and used in the LoD analysis. **a**. 5 min assay time. **b**. 10 min assay time. **c**. 15 min assay time. **d**. 20 min assay time.
